## Supplementary Information for "Premature commitment to uncertain beliefs during human NMDA receptor hypofunction"

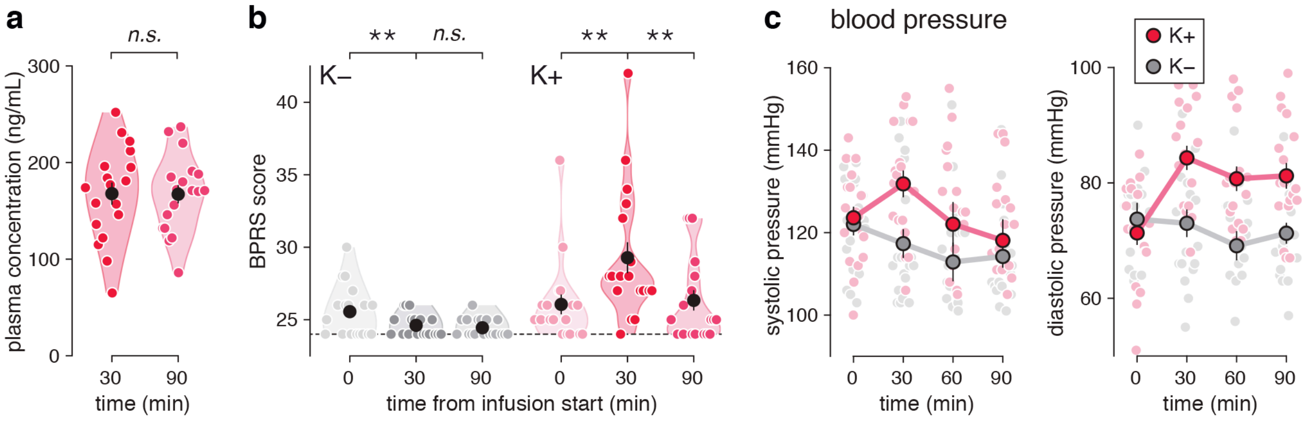


**Supplementary Fig. 1 | Pharmacological protocol and effects. a**, Plasma concentration of ketamine before ($t$ = 30 min) and after ($t \approx$ 90 min) task execution. Black dots and error bars indicate group-level means ± s.e.m., whereas colored dots indicate participant-level data. The plasma concentration of ketamine is stable and close to target during task execution. **b**, Scores on the Brief Psychiatric Rating Scale (BPRS). The BPRS score increases under ketamine, with only four BPRS items (out of 24) showing significant increases: elated mood, excitement, distractibility and unusual thought content. **c**, Side effect of ketamine on blood pressure (left: systolic pressure; right: diastolic pressure). Blood pressure shows a previously reported mild elevation under ketamine. Large dots and error bars indicate group-level means ± s.e.m., whereas small dots indicate participant-level data. Two stars indicate a significant effect at $p < 0.01$, *n.s.* a non-significant effect.

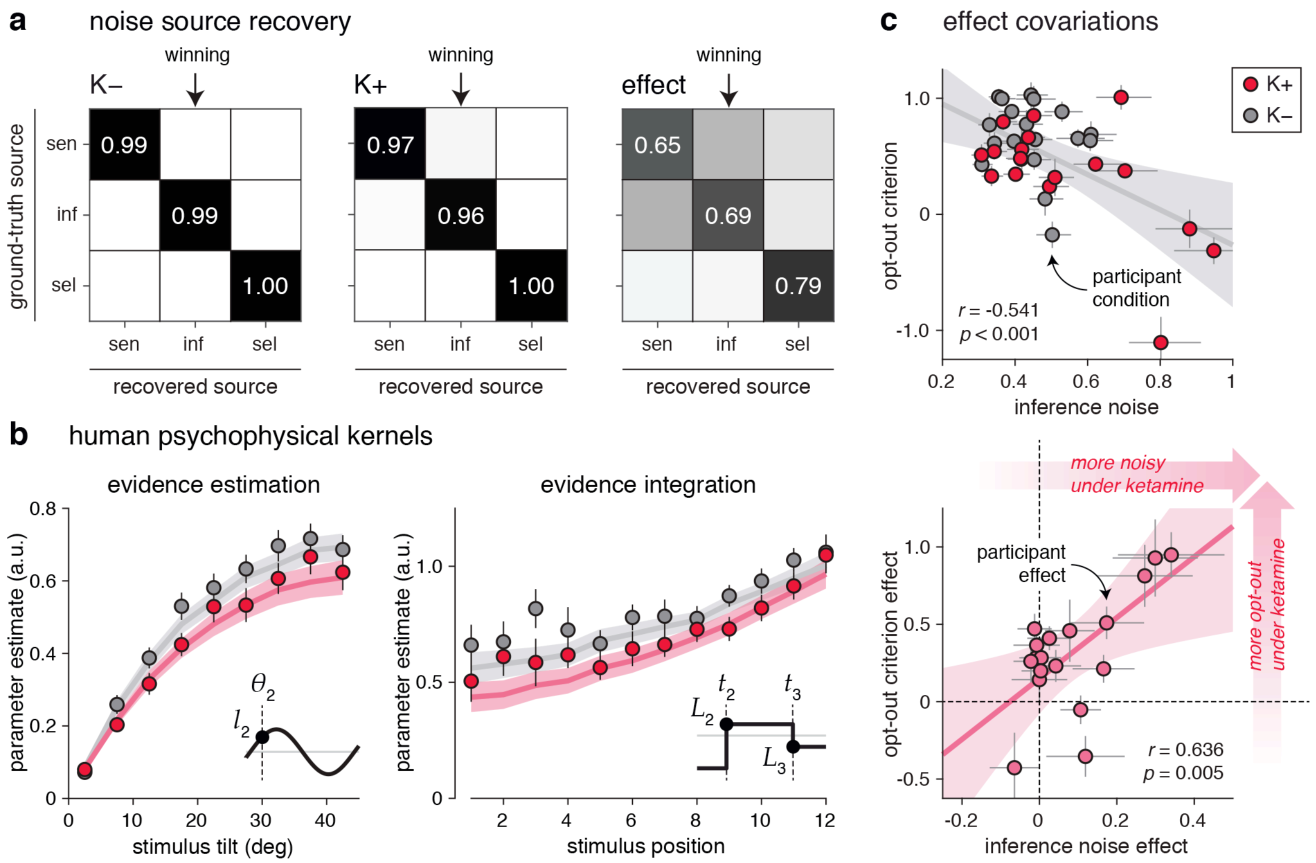


**Supplementary Fig. 2 | Suboptimal inference model validation. a**, Noise source recovery procedure for the placebo condition (left), the ketamine condition (middle), and the difference between conditions (right). Confusion matrices show the probability of each recovered noise source (columns) being the ground-truth source (rows), based on model simulations using group-level means as parameter values. All three noise sources can be correctly identified. The confusion matrix for the difference between conditions is mainly diagonal, but increased sensory noise can be confused with increased inference noise (confusion $p$ = 0.308) and vice versa. **b**, Human psychophysical kernels with respect to stimulus tilt from the closest category boundary (evidence estimation) and stimulus position (evidence integration). Dots indicate observations (group-level means ± s.e.m.), whereas lines indicate best-fitting predictions. Participants estimate the evidence equally well, and exhibit moderate evidence integration leaks in the two conditions. **c**, Covariation between psychometric parameters and effects of ketamine. Top: negative correlation between inference noise (x-axis) and opt-out criterion (y-axis). Each dot corresponds to one condition (placebo: gray, ketamine: red) of a single participant. Participants making more inference errors use a lower opt-out criterion. Right: positive correlation between the effects of ketamine on inference noise (x-axis, positive means more noise under ketamine) and on the opt-out criterion (y-axis, positive means more frequent opt-out under ketamine). Each dot corresponds to the difference between conditions (ketamine *minus* placebo) for a single participant. Large increases in inference noise are associated with large decreases in the opt-out criterion. Error bars correspond to marginal s.d. of best-fitting parameter values. Lines correspond to best-fitting regression lines, shaded error bars to their 95% confidence intervals.

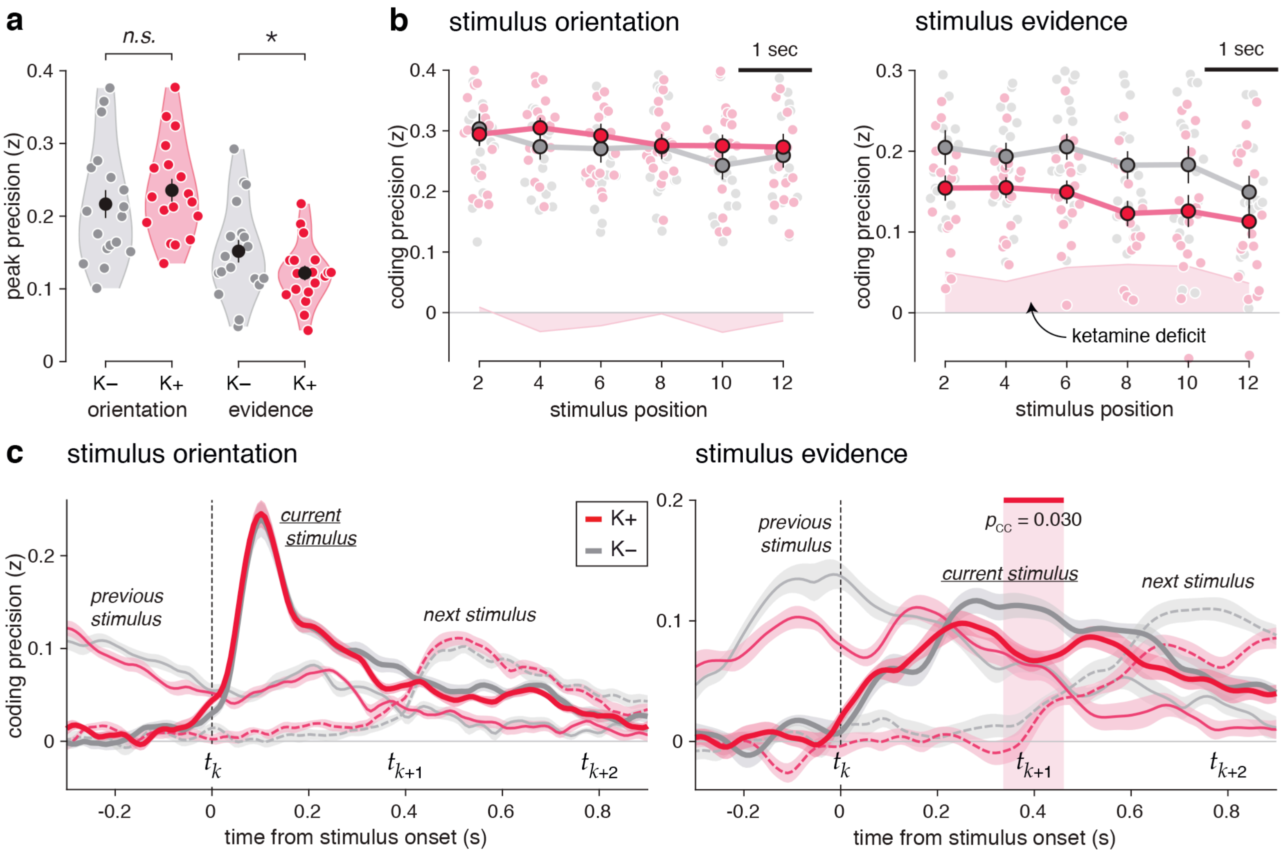


**Supplementary Fig. 3 | Temporal properties of neural coding. a**, Peak precision of neural coding for stimulus orientation (left) and stimulus evidence (right). The peak precision of orientation coding does not differ between conditions, whereas the peak precision of evidence coding is significantly reduced under ketamine. **b**, Coding precision as a function of stimulus position in the sequence. Left: neural coding of stimulus orientation at 50-150 ms following stimulus onset. Right: neural coding of stimulus evidence at 250-450 ms following stimulus onset. The red-shaded area indicates the coding deficit under ketamine. The neural coding of stimulus evidence is degraded across all stimulus positions under ketamine. **c**, Overlapping neural codes of successive stimuli (left: stimulus orientation; right: stimulus evidence). Simultaneous decoding of previous ($k$−1, thin solid lines), current ($k$, thick solid lines) and next ($k$+1, thin dashed lines) stimuli from EEG signals aligned to the onset of stimulus $k$. Lines and shaded error bars indicate group-level means ± s.e.m. The neural coding of orientation overlaps only slightly across successive stimuli, whereas the neural coding of evidence overlaps strongly across successive stimuli. The red-shaded area indicates the significant decrease in coding precision under ketamine for stimulus $k$, accounting for the simultaneous neural coding of previous and next stimuli. One star indicates a significant effect at $p < 0.05$, *n.s.* a non-significant effect.

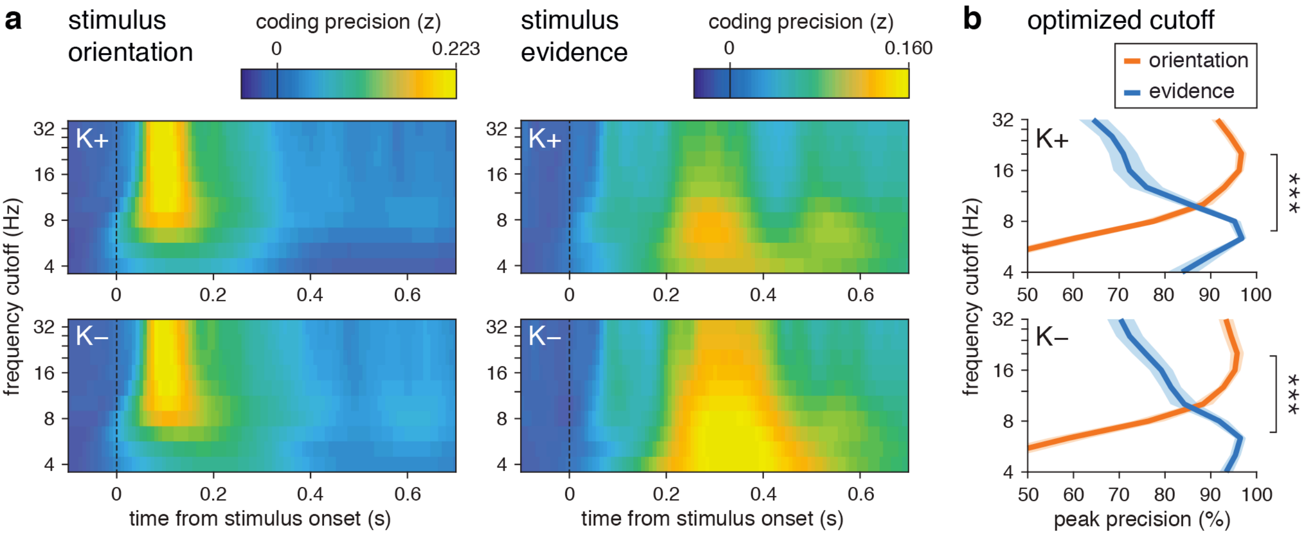


**Supplementary Fig. 4 | Spectral properties of neural coding. a**, Effect of frequency cutoff (y-axis) on the neural coding of stimulus orientation (left) and stimulus evidence (right) in the ketamine (top) and placebo (bottom) conditions. The neural coding of stimulus orientation is supported by spectral content above 8 Hz, whereas the neural coding of stimulus evidence is supported by spectral content below 8 Hz. **b**, Optimized frequency cutoff (y-axis) for the neural coding of stimulus orientation and evidence in the ketamine (top) and placebo (bottom) conditions. Lines and error bars correspond to group-level means ± s.e.m. The optimal frequency cutoff (for which the peak coding precision is maximal) is slightly above 16 Hz for stimulus orientation, and slightly below 8 Hz for stimulus evidence. This difference in optimal frequency cutoff is highly significant in both conditions. Three stars indicate a significant effect at $p < 0.001$.

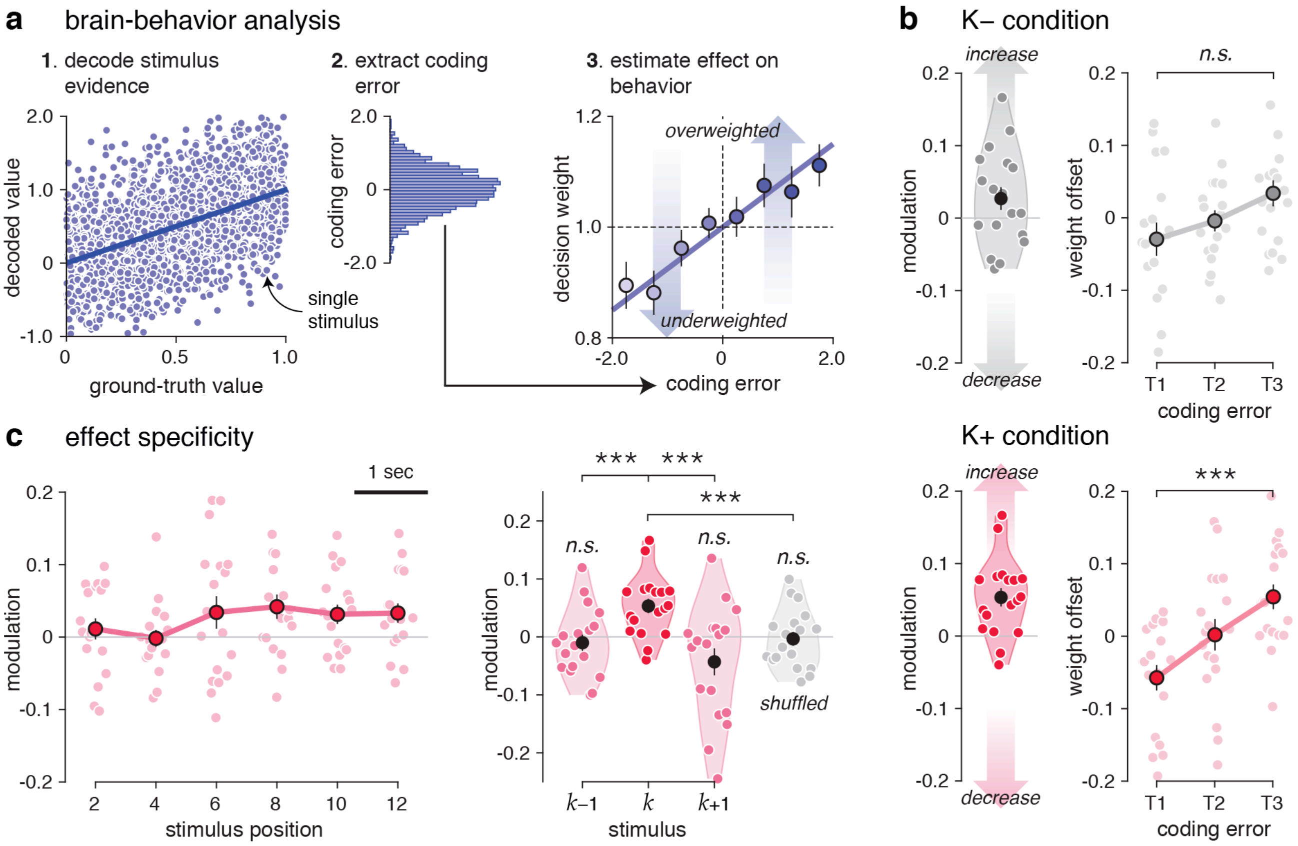


**Supplementary Fig. 5 | Relation between neural coding and decision weight. a**, Description of the 'brain-behavior' analytic pipeline. Left: stimulus evidence is decoded from EEG signals (step 1). Center: evidence coding errors are computed as the difference between the decoded stimulus evidence $\hat{x}$ and the ground-truth stimulus evidence $x$ (step 2). Positive coding errors indicate overestimated evidence, whereas negative coding errors indicate underestimated evidence. Right: the effect of evidence coding errors (x-axis) on decision weight (y-axis) is estimated by entering evidence coding errors as a multiplicative modulation factor for the ground-truth evidence provided by the same stimulus in the suboptimal inference model (step 3). **b**, Relation between evidence coding errors at 250-450 ms following stimulus onset and decision weight (top: placebo; bottom: ketamine). Left: modulation parameter estimate (positive means that overestimated evidence is associated with increased decision weight). Right: decision weight offset (y-axis) from its average value (positive means overweighting, negative means underweighting) as a function of evidence coding errors (x-axis) binned into terciles. Evidence coding errors correlate positively with decision weight under ketamine. **c**, Effect specificity. Left: modulation parameter estimate as a function of stimulus position in the sequence under ketamine. The relation between evidence coding errors and decision weight emerges at the same time as the coding unbalance. Right: lagged modulation kernel, corresponding to the effects of evidence coding errors associated with stimuli $k-1$, $k$ and $k+1$ on the decision weight $w_k$ assigned to stimulus $k$. Only coding errors associated with stimulus $k$ co-vary with the decision weight assigned to stimulus $k$ in the subsequent decision. Shuffling evidence coding errors across stimuli (in gray) abolishes the effect of evidence coding errors on decision weight. Three stars indicate a significant effect at $p < 0.001$, *n.s.* a non-significant effect.

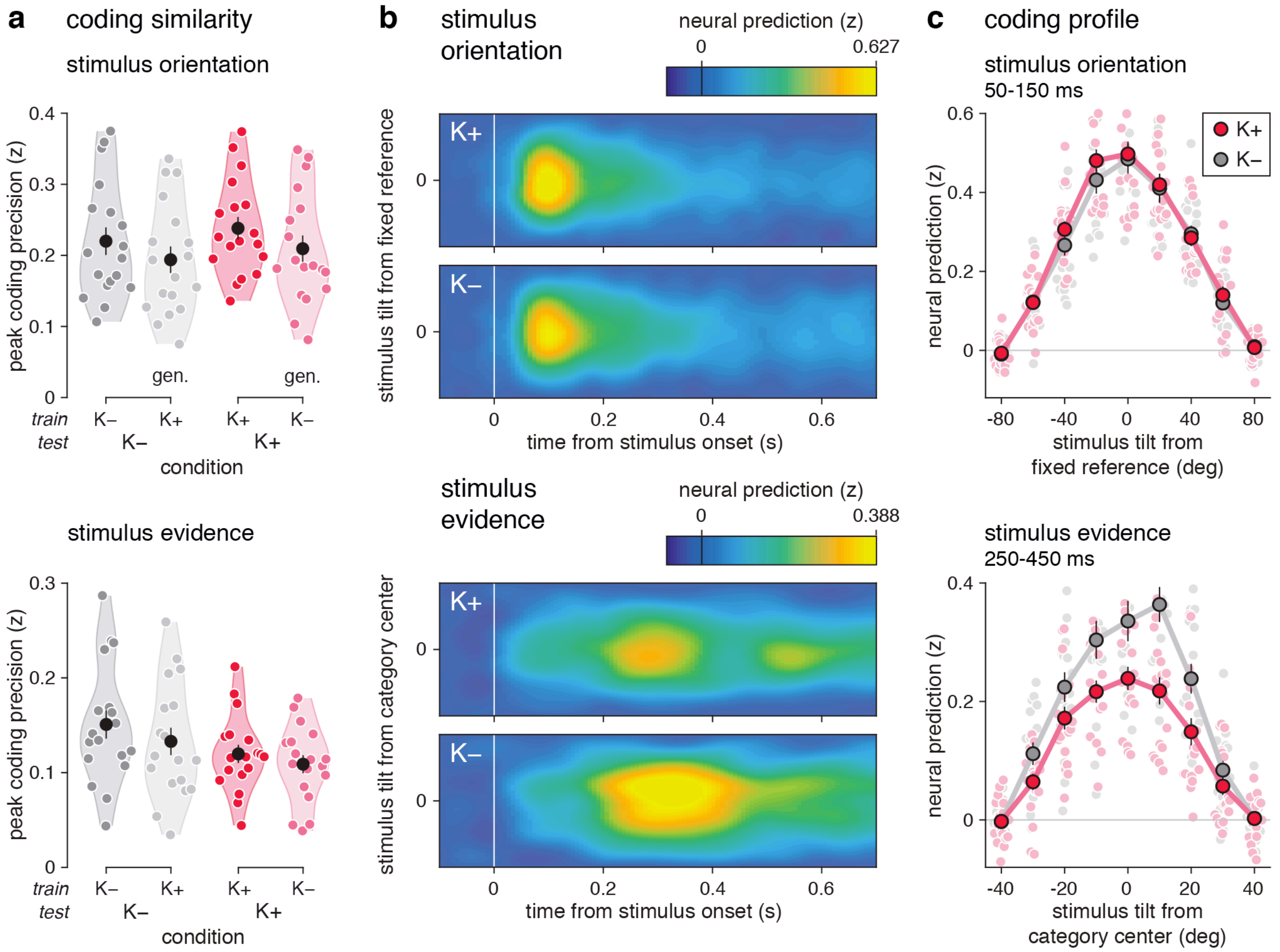


**Supplementary Fig. 6 | Shared neural codes across conditions. a**, Cross-condition generalization analysis of neural codes (top: stimulus orientation; bottom: stimulus evidence) Using spatial patterns trained in one condition to compute cross-validated neural predictions in the other condition has only a subtle negative impact on peak coding precision. This pattern suggest a high degree of coding similarity in the neural codes of each stimulus characteristic between conditions. **b**, Time course of the gain of neural coding (top: stimulus orientation; bottom: stimulus evidence). Coding profile of neural predictions over time (x-axis) as a function of stimulus tilt (y-axis) from a fixed reference (stimulus orientation) or from the closest category center – which changes from stimulus to stimulus and from trial to trial (stimulus evidence). Neural predictions are derived using the same spatial patterns for decoding pooled across conditions. **c**, Coding profile of neural predictions (top: stimulus orientation; bottom: stimulus evidence) around their peak. Stimulus orientation is processed with equal neural gain in the two conditions, whereas stimulus evidence is processed with lower neural gain under ketamine at 250-450 ms following stimulus onset.

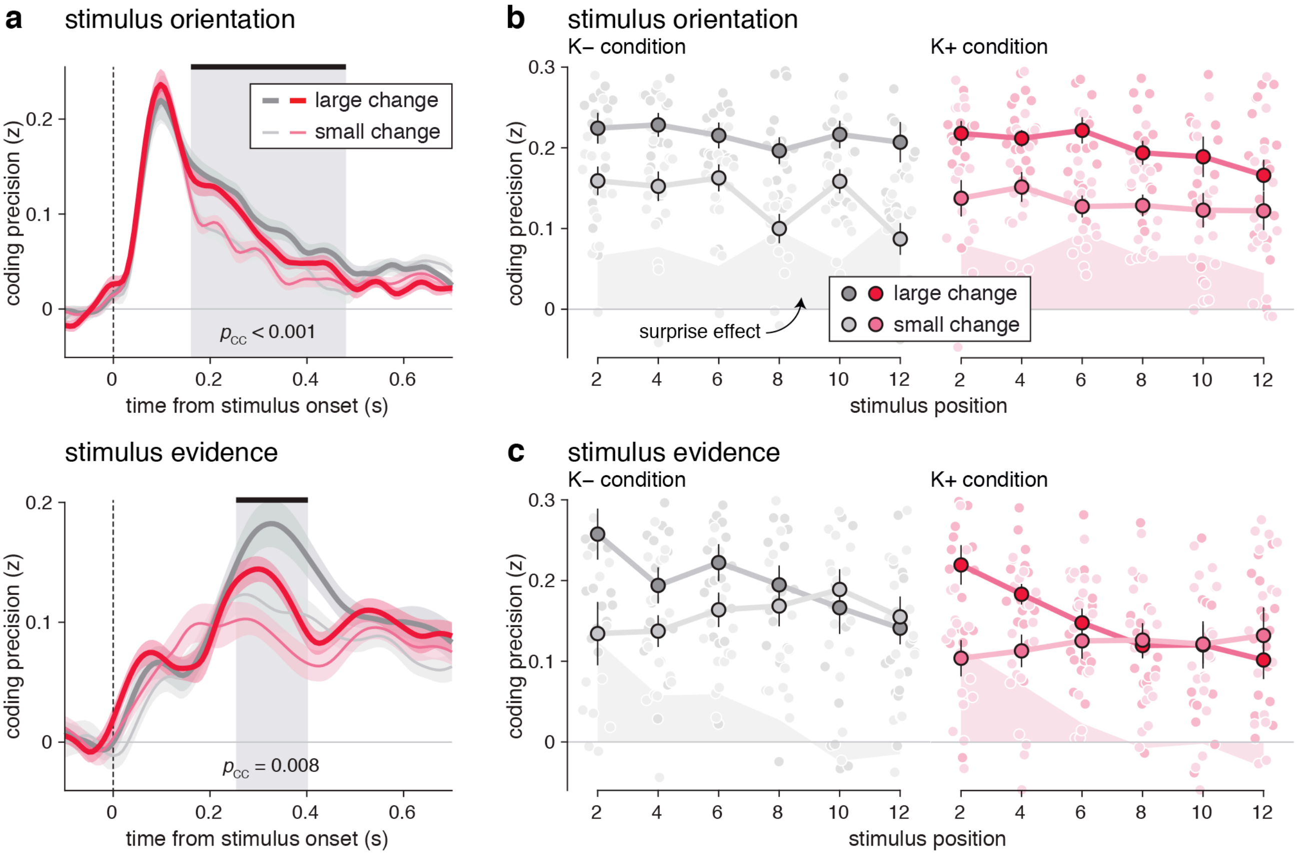


**Supplementary Fig. 7 | Effects of stimulus change on neural coding. a**, Effects of stimulus change on neural coding (top: stimulus orientation; bottom: stimulus evidence). Thick lines correspond to stimuli associated with large tilt (more than 45 degrees) from their predecessor in the sequence, whereas thin lines correspond to stimuli associated with small tilt from their predecessor (less than 45 degrees). The neural coding of stimulus orientation shows a strong and sustained 'surprise' effect across conditions at 160-480 ms following stimulus onset (indicated by the gray-shaded area). The neural coding of stimulus evidence also shows a 'surprise' effect across conditions at 250-400 ms following stimulus onset. **b**, Effect of stimulus change on orientation coding as a function of stimulus position (left: placebo; right: ketamine). Shaded areas indicate the strength of surprise effect (i.e., the difference in coding precision between stimuli associated with large tilt and those associated with small tilt from their predecessor). The surprise effect on orientation coding is sustained over time in both conditions. **c**, Effect of stimulus change on evidence coding as a function of stimulus position (left: placebo; right: ketamine). The surprise effect on evidence coding fades over the course of the sequence in both conditions.

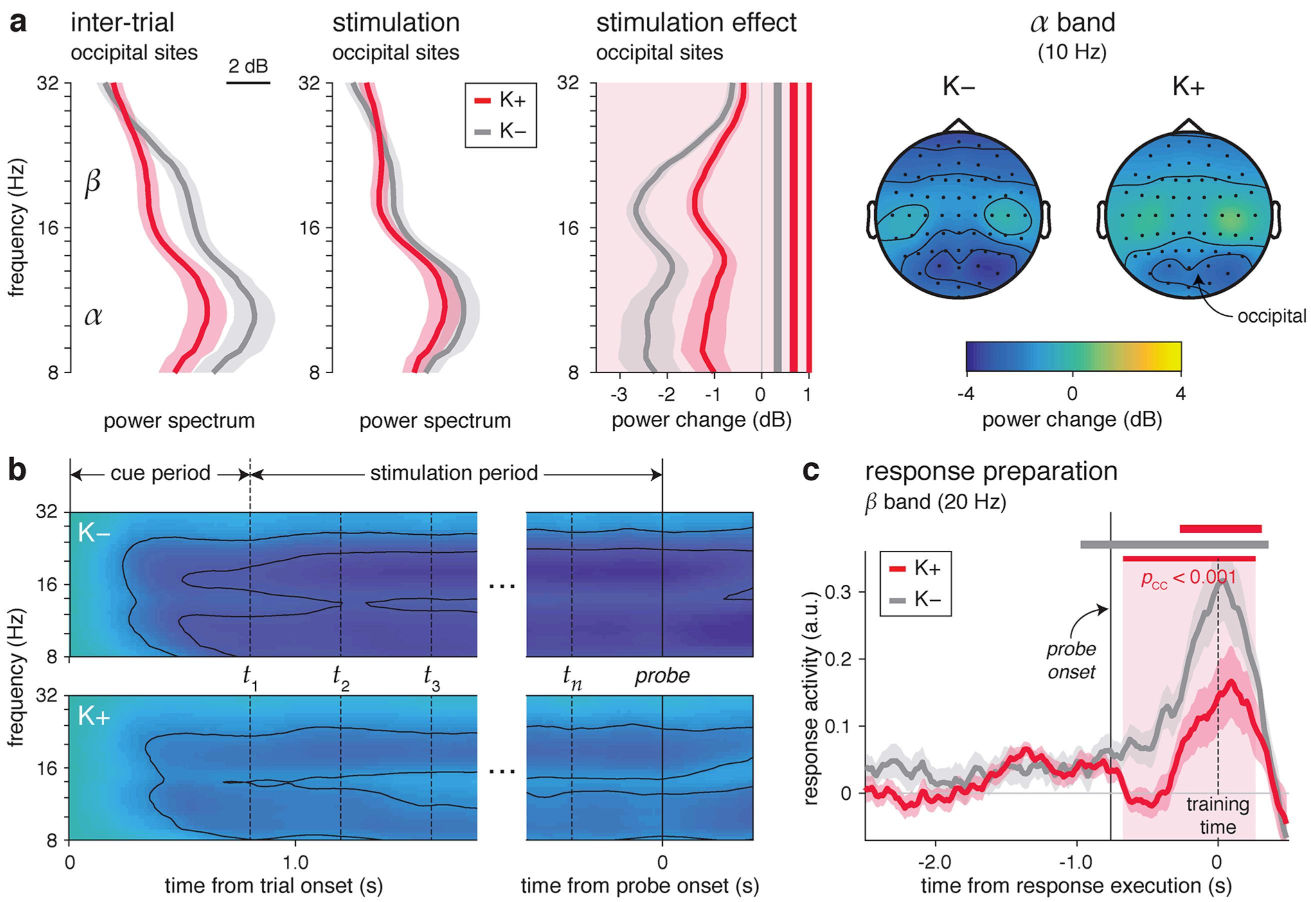


**Supplementary Fig. 8 | Spectral alterations under ketamine.** **a**, Suppression of alpha power at occipital channels. Left: power spectrum (x-axis) between 8 and 32 Hz at occipital channels during the inter-trial interval (left), visual stimulation (middle), and the stimulation effect (stimulation *minus* inter-trial, right). Inter-trial power is decreased under ketamine across alpha and beta bands, but less so during visual stimulation, corresponding to a dampened suppression of alpha and beta power under ketamine. The red-shaded area indicates frequencies showing a significant difference in power suppression between conditions. Right: spatial topography of the suppression of alpha power (10 Hz) during visual stimulation. The suppression of alpha power peaks at occipital channels overlying human visual cortex, and is dampened under ketamine. **b**, Time-frequency diagram of power suppression (top: placebo; bottom: ketamine) at occipital channels over time (x-axis) and frequencies (y-axis). The dampened suppression of alpha and beta power under ketamine is visible throughout each trial, from the cue period (before visual stimulation) until the response period (following the 'go' signal). Contours delineate power decreases of 1 and 2 dB. **c**, Time course of beta power (20 Hz) projected on the response-predictive axis (response activity) in the last 2.5 s preceding response execution. Response activity in beta power is less accurate under ketamine during response execution (indicated by the red-shaded area). Shaded error bars indicate s.e.m.

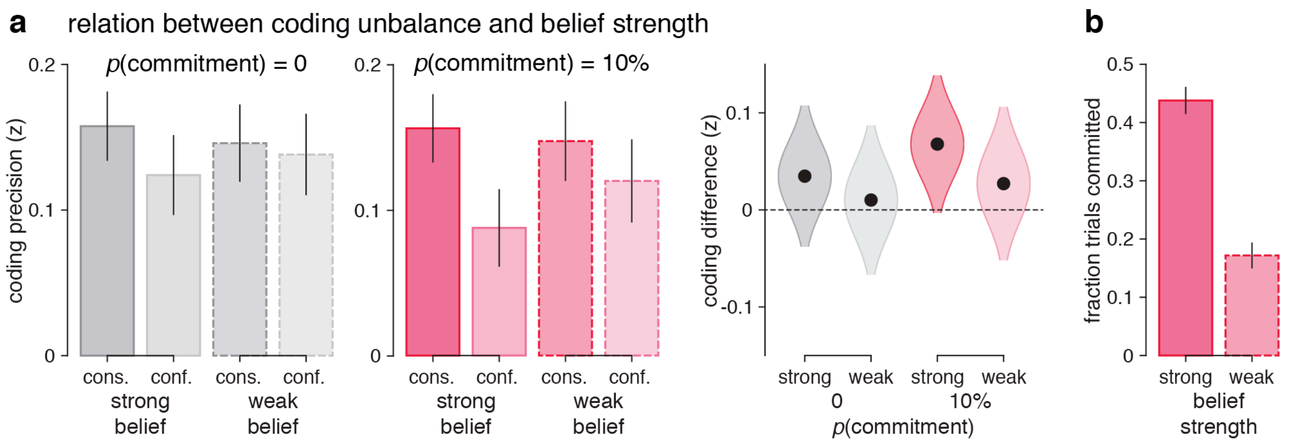


**Supplementary Fig. 9 | Relation between premature commitments and belief strength. a**, Predicted relation between coding unbalance and belief strength, obtained using simulations of suboptimal cognitive inference without premature commitment (gray bars and violins, $p$ = 0) or with premature commitments (red bars and violins, $p$ = 10%). Left: coding precision of consistent and conflicting evidence as a function of the amount of accumulated evidence at the end of the trial (strong vs. weak belief), and the presence of premature commitments ($p$ = 0 vs. $p$ = 10%). Right: coding unbalance (i.e., the difference in coding precision between consistent and conflicting evidence) as a function of the same two factors. Coding unbalance is more pronounced in trials ending with a larger amount of accumulated evidence, whether premature commitments occur or not. Bars (or dots) and error bars (or violins) indicate group-level means ± s.e.m. of coding precision estimates across simulations. **b**, Fraction of simulated trials with premature commitment as a function of the amount of accumulated evidence at the end of the trial. Premature commitments are substantially more likely to have occurred in trials ending with a larger than average amount of accumulated evidence (strong belief). Bars and error bars indicate group-level means ± s.e.m for simulations with $p$(commitment) = 10%.

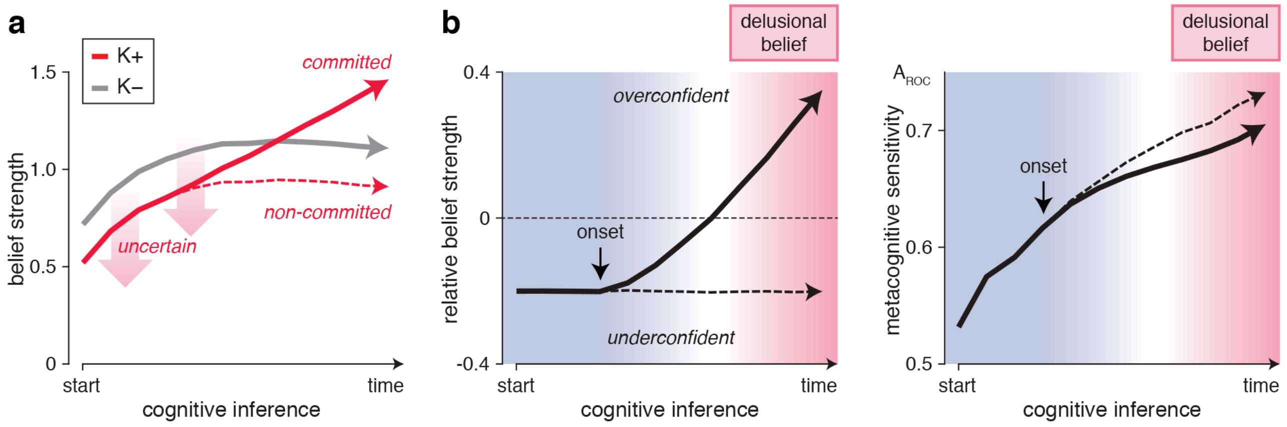


**Supplementary Fig. 10 | Proposed account of the pathogenesis of psychosis. a**, Time course of belief strength in a noisy and leaky cognitive inference model, for 'regular' trials without premature commitment (in gray) and 'altered' trials with increased decision uncertainty and premature commitments (in red). The dashed red curve corresponds to altered trials where no premature commitment has occurred, whereas the solid red curve corresponds to altered trials where a premature commitment has occurred. The occurrence of premature commitments in altered trials compensates for the increased decision uncertainty due to the selective integration of consistent evidence, and the discarding of conflicting evidence. **b**, Transition from uncertain to delusional beliefs in a premature commitment model of cognitive inference. Left: time course of relative belief strength (i.e., the difference in belief strength between altered and regular trials). The early underconfidence (left) found in altered trials gets compensated by the selective integration of consistent evidence in trials where a premature commitment occurs (solid curve), and triggers late overconfidence in inferred beliefs (right). This diachronic pattern is robust to changes in parameter values. Right: time course of metacognitive sensitivity (i.e., the standardized difference in belief strength between correct decisions and errors). Premature commitments progressively degrade metacognitive sensitivity due to the unbalanced integration of evidence, resulting in delusional beliefs (stronger and less accurate than normal).
